# Comprehensive detection of structural variation and transposable element differences between wild type laboratory lineages of *C. elegans*

**DOI:** 10.1101/2023.01.13.523974

**Authors:** Zachary D. Bush, Alice F. S. Naftaly, Devin Dinwiddie, Cora Albers, Kenneth J. Hillers, Diana E. Libuda

**Author notes:** Corresponding author <u>Corresponding Author and Lead Contact Information:</u> Diana E. Libuda, PhD University of Oregon Institute of Molecular Biology 1229 Franklin Blvd Eugene, OR 97403.

## Abstract

Genomic structural variations (SVs) and transposable elements (TEs) can be significant contributors to genome evolution, altered gene expression, and risk of genetic diseases. Recent advancements in long-read sequencing have greatly improved the quality of *de novo* genome assemblies and enhanced the detection of sequence variants at the scale of hundreds or thousands of bases. Comparisons between two diverged wild isolates of *Caenorhabditis elegans*, the Bristol and Hawaiian strains, have been widely utilized in the analysis of small genetic variations. Genetic drift, including SVs and rearrangements of repeated sequences such as TEs, can occur over time from long-term maintenance of wild type isolates within the laboratory. To comprehensively detect both large and small structural variations as well as TEs due to genetic drift, we generated *de novo* genome assemblies and annotations for each strain from our lab collection using both long- and short-read sequencing and compared our assemblies and annotations with that of other lab wild type strains. Within our lab assemblies, we annotate over 3.1Mb of sequence divergence between the Bristol and Hawaiian isolates: 337,584 SNPs, 94,503 small insertion-deletions (<50bp), and 4,334 structural variations (>50bp). Further, we define the location and movement of specific DNA TEs between N2 Bristol and CB4856 Hawaiian wild type isolates. Specifically, we find the N2 Bristol genome has 20.6% more TEs from the *Tc1/mariner* family than the CB4856 Hawaiian genome. Moreover, we identified Zator elements as the most abundant and mobile TE family in the genome. Using specific TE sequences with unique SNPs, we also identify 38 TEs that moved intrachromosomally and 9 TEs that moved interchromosomally between the N2 Bristol and CB4856 Hawaiian genomes. By comparing the *de novo* genome assembly of our lab collection Bristol isolate to the VC2010 Bristol assembly, we also reveal that lab lineages display over 2 Mb of total variation: 1,162 SNPs, 1,528 indels, and 897 SVs with 95% of the variation due to SVs. Overall, our work demonstrates the unique contribution of SVs and TEs to variation and genetic drift between wild type laboratory strains assumed to be isogenic despite growing evidence of genetic drift and phenotypic variation.

**Author Summary:** For multiple model organisms, propagation of wild type strains in independent labs can lead to multiple phenotypic differences over time. To assess recombination, map mutations, and understand genomic changes during speciation, *Caenorhabditis elegans* researchers primarily use the wild type isolates Bristol and Hawaiian. Here, we map structural variations, transposable elements, and sequence divergence between the Bristol and Hawaiian natural isolates and between genomes of different lab lineages of these same strains.

## Introduction

Genomic variants, through mutation and recombination, in individuals and genetic drift in populations underly the core process of evolution. Functional characterization of sequence variants guides our understanding of phenotypic variances within species while also being critical to identifying heritable disease-causing mutations (Haraksingh and Snyder 2013). Genomic variation has been reported at multiple scales, from single nucleotide polymorphisms (SNPs) to short insertions/deletions (indels) to much larger structural variants (SVs). SVs are defined as insertions, deletions, or chromosomal rearrangements at least 50bp in length. SVs can cause loss of function mutations through large gene deletions or alter gene expression by disrupting spatial interactions between regulatory sequences (Stranger et al. 2007; Hurles, Dermitzakis, and Tyler-Smith 2008). Accurate detection of both sequence variants and chromosome rearrangements is critical for understanding how genomic variation may contribute to phenotypic plasticity in individuals and populations of the same species.

Transposable elements (TEs) are a class of repetitive DNA sequences capable of moving to new locations in the genome. TE mobility is a source of genomic structural variation that can also alter gene expression (Girard and Freeling 1999; Slotkin and Martienssen 2007) and drive, sometimes rapid, evolutionary changes within species (Van’t Hof et al. 2016; Feschotte and Pritham 2007). Notably, transposons account for a significant fraction of the total DNA sequence in many eukaryotic species (Chalopin et al. 2015; Gilbert, Peccoud, and Cordaux 2021), which provides many opportunities for TE-driven structural rearrangements. The *Tc1/mariner* family of DNA transposons is one of the most abundant TEs across species (Eide and Anderson 1985; Plasterk, Izsvák, and Ivics 1999), and early studies in *C. elegans* found it to be one of the few mobile transposons observed under laboratory conditions (Fischer, Wienholds, and Plasterk 2003). To repress or limit transposon mobilization, transposon silencing is tightly regulated through multiple mechanisms including chromatin modification and RNA interference (Sijen and Plasterk 2003; H.-C. Lee et al. 2012). Despite their ubiquity and impact on genomic architecture, the comprehensive annotation and inclusion of TEs in comparative genomic analyses has been challenging. Many studies have incompletely characterized the genomic distribution of TEs because older, short-read based genome assemblies could not accurately map the full content of repetitive sequences. Further, programs that automatically detect TEs based on sequence homology and conserved sequence elements rely heavily on libraries of older reference sequences that may predate the discovery of TE fragments and newer TE families. As new families of transposable elements are discovered (Bao et al. 2009) along with new technology that aids their annotation and tracking (Riehl et al. 2022), determining the genomic composition and mobility of new TEs will enable our understanding of their role in genome evolution and genome integrity.

Foundational research on genomic variation has utilized next generation short-read sequencing, long-read sequencing, and the direct comparison of reference genome assemblies to identify genomic variants (Mahmoud et al. 2019; Lappalainen et al. 2019). SNPs and indels, ranging in size from 1 bp to 50bp, can be identified with high confidence using short sequencing reads that are 100-150bp (Muzzey, Evans, and Lieber 2015). In contrast, SVs are challenging to annotate using short-read sequencing because the sequencing reads are often smaller than the size of an SV (Sudmant et al. 2015; Mahmoud et al. 2019; Lesack et al. 2022). Similarly, the highly repetitive sequences of TEs present significant challenges to mapping and annotation with traditional short read sequencing methods. With the advent of higher quality long-read sequencing technologies which generate ∼10kb-30kb reads with lower genomic coverage, the accurate annotation of large regions of genomic variation such as SVs and transposable elements has become easier (Sakamoto et al. 2021). New tools to identify SVs via assembly-to-assembly alignments (Delcher et al. 1999; Nattestad and Schatz 2016; Li 2018; Goel et al. 2019) are not constrained by read-length to identify SVs and depend on high-quality reference assemblies. Thus, a high-quality reference genome assembly is a critical resource for any model organism. Methods of variant detection that leverage a combined utilization of short- and long-read sequencing can provide more accurate reference sequences to fully address undiscovered genomic variations previously not detected by short-read sequencing alone.

*Caenorhabditis elegans* was the first multicellular organism to have its genome fully sequenced (C. elegans Sequencing Consortium 1998) and has been exploited to pioneer many comparative genomic studies. To understand how genetic variation influences phenotypic differences and genomic processes within species, *C. elegans* researchers primarily utilize two highly diverged wild type strains estimated to have diverged 30,000-50,000 generations ago (Thomas et al. 2015): N2 (isolated in Bristol, England) and CB4856 (isolated in Maui, Hawaii) (Nicholas, Dougherty, and Hansen 1959; Sulston and Brenner 1974; Hodgkin and Doniach 1997; Crombie et al. 2019). Earlier comparisons of the Bristol and Hawaiian lineages were critical for studying genetic variation, gene families, and evolution of genome structures (Koch et al. 2000; Wicks et al. 2001; Stewart et al. 2005; Maydan et al. 2010). The *C. elegans* genome, comprised of 5 autosomes and the X chromosome, displays a nonuniform distribution of sequence variation when comparing the genomes of wild isolates. Although a large amount of sequence divergence was previously found between the N2 Bristol and CB4856 Hawaiian lineages (Thompson et al. 2015; Andersen et al. 2012), the increased quality of reference genomes, sequencing technology, and variant detection methods enables the identification of additional variations (in particular large structural variations) that previously went undetected in these *C. elegans* genomes.

Recently, Bristol and Hawaiian genomes were reassembled *de novo* using a combination of short-read Illumina sequencing as well as long-read sequencing from PacBio and Oxford Nanopore platforms (Yoshimura et al. 2019; Kim et al. 2019). Compared to the previous short-read based assemblies of N2 Bristol, the new assembly of N2 Bristol, called VC2010, identified 53 more predicted genes, 1.8Mb of additional sequence, and eliminated 98% of existing gaps in the N2 Bristol genome. Thus, the VC2010 Bristol genome very likely better represents the genome of Bristol *C. elegans* currently used in laboratories worldwide (Yoshimura et al. 2019). The first CB4856 Hawaiian genome assembly was completed in 2015 by iteratively correcting the pre-existing N2 Bristol reference assembly (*C. elegans* Sequencing Consortium 1998) with short-read sequencing data (Thompson et al. 2015). This study identified 327,050 single-nucleotide polymorphisms (SNPs) and nearly 80,000 indels relative to N2; a marked increase relative to previous comparisons, which had identified 6,000-17,000 SNPs and small indels (Wicks et al. 2001; Swan et al. 2002) between N2 Bristol and CB4856 Hawaiian. Due to the size of the short read sequences employed in the analysis, the iterative correction method used to assemble the CB4856 Hawaiian genome may not have detected all structural rearrangements and repetitive sequences. In 2019, the first *de novo* CB4856 Hawaiian assembly from long-read sequencing extended the length of the Hawaiian genome, and was further able to characterize over 3,000 previously uncharacterized SVs (Kim et al. 2019). Thus, combining long-read and short-read sequencing in *de novo* genome assembly not only extended the known length of both the N2 Bristol and CB4856 Hawaiian isolate genomes, but broadened our understanding of how much genomic variation exists between these wild-type strains.

Many *C. elegans* research labs utilize N2 Bristol and CB4856 Hawaiian as standard wild type strains, but long-term passaging in each lab may lead to the accumulation of many smaller sequence variants and large genomic structural variations. Early assessments of laboratory lineages of the N2 Bristol strain, for example, identified many duplications ranging in size from 200bp to 108kb, with some affecting as many as 26 genes (Vergara et al. 2009). To determine the extent of genetic variation between our laboratory lineages of N2 Bristol and CB4856 Hawaiian, we generated two high-quality reference assemblies for the N2 and CB4856 strains used in our laboratory to compare to that of other high-quality reference genomes for N2 and CB4856. By leveraging recent technological advancements in sequencing and variant detection, we provide a comprehensive annotation of SNPs, indels, structural variations, and transposable elements between our lineages of the Bristol and Hawaiian strains. From our comprehensive mapping of TEs in our reference genomes, we report Zator elements to be the most abundant and mobile TE family in the *C. elegans* genome. Further, by comparing our assembled genomes to recently published VC2010 Bristol and CB4856 Hawaiian long-read assemblies (Yoshimura et al. 2019; C. Kim et al. 2019), we identified SNPs, indels, and SVs unique to different lab wild type strains. These variations were enriched in intergenic regions of the *C. elegans* genome, suggesting that variations in regulatory sequences and other non-coding regions may underlie the phenotypic variances previously observed between laboratory strains. Taken together, our systematic analysis of genetic variation between natural and laboratory wild type isolates highlights the impact of large structural variants, TE composition, and other chromosomal rearrangements accumulating in the genomes of laboratory model organisms.

## Results

### *De novo* genome assembly using combined long and short-read sequencing produces high quality genomes

To perform systematic comparisons of multiple wild type genomes from different laboratory isogenic strains, we generated *de novo* assemblies of N2 Bristol and CB4856 Hawaiian. The N2 Bristol genome was assembled from PacBio long-reads with 136x coverage producing 121 contigs and a 100.4Mb genome (Figure 1A) The CB4856 Hawaiian genome was generated from PacBio long-reads with 132x coverage from 169 contigs to give a 98.8Mb assembly (Figure 1B). These long-read assemblies were then supplemented with Illumina paired end short-reads with a sequencing depth of 540x and 628x for N2 Bristol and CB4856 Hawaiian respectively (Figure 1A-B).

**Figure 1.**
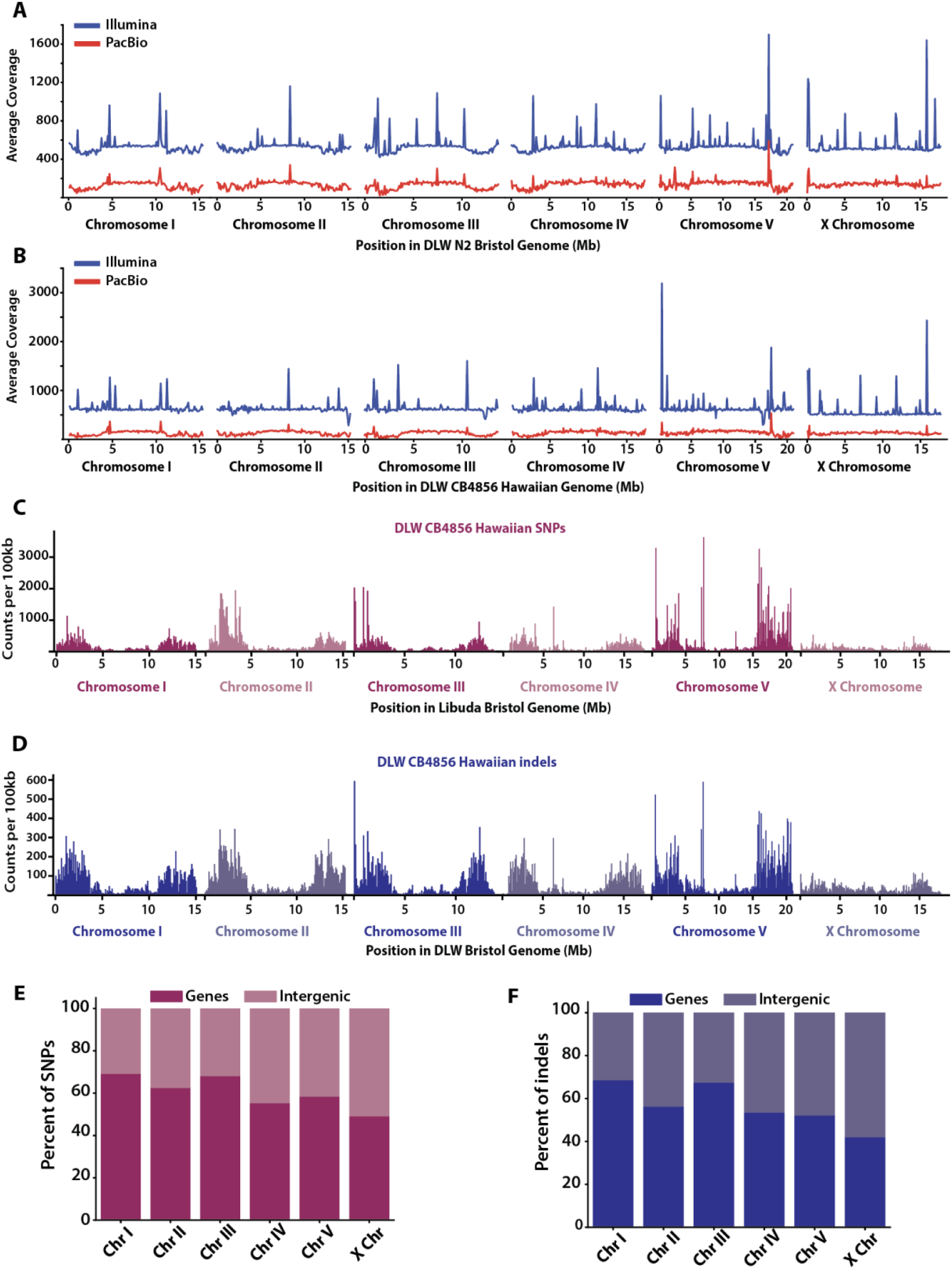
Genomic distribution of SNPs and indels between the DLW N2 Bristol and DLW CB4856 Hawaiian genomes. **(A)** Line plots showing the average sequencing coverage in 100kb bins across each chromosome in the DLW N2 Bristol genome. **(B)** Line plots showing the average sequencing coverage in 100kb bins across each chromosome in the DLW CB4856 Hawaiian genome. For each plot in A and B, the coverage for Illumina short-read sequencing is shown in blue, and sequencing coverage for PacBio long-reads is shown in red. **(C)** Histograms depicting the distribution of CB4856 Hawaiian SNPs across each DLW N2 Bristol chromosome in 100kb bins. **(D)** Histograms of the distributions of CB4856 Hawaiian indels across each DLW N2 Bristol chromosome in 100kb bins. **(E)** The proportion of SNPs that overlap with remapped gene annotations versus intergenic regions in the DLW N2 Bristol genome. **(F)** The proportion of indels that overlap with gene versus intergenic regions in the Bristol genome.

To assess the quality of our reference genomes, we examined assembly-to-assembly alignments and the orthologous gene content for each assembly. A strong assembly would show a similar proportion of aligned bases and a high degree of synteny when comparing across assemblies. In concordance with comparisons in earlier studies, 99.2% of bases across our N2 Bristol and CB4856 Hawaiian assembled genomes were aligned (Kim et al. 2019), and more than 92.2% of bases within alignments were syntenic (Table 1).

**Table 1.**
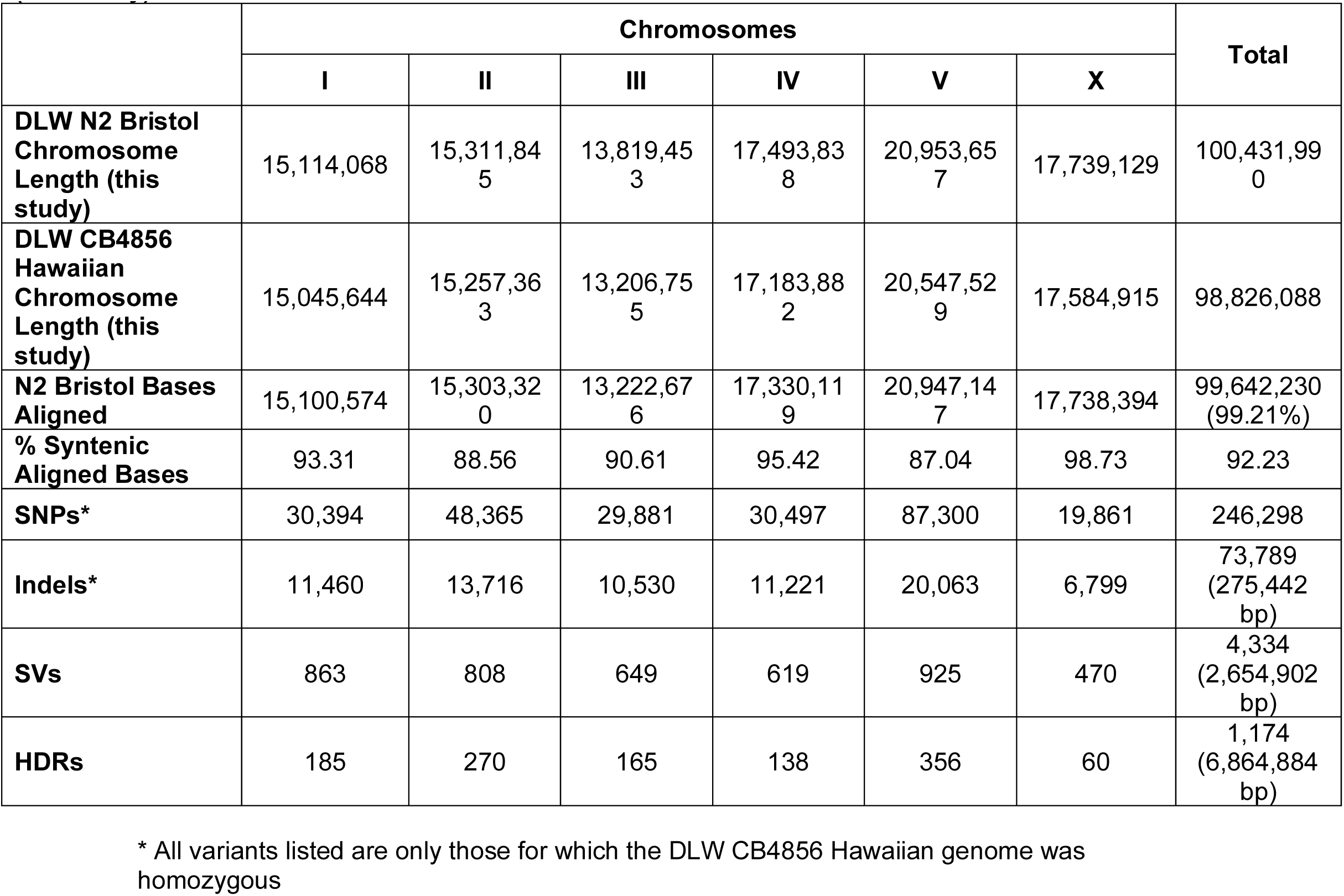
Comparisons between the DLW N2 Bristol genome (this study) and DLW CB4856 Hawaiian genome dy)

|  | Chromosomes |  |  |  |  |  | Total |
| --- | --- | --- | --- | --- | --- | --- | --- |
|  | I | II | III | IV | V | X |  |
| **DLW N2 Bristol Chromosome Length (this study)** | 15,114,068 | 15,311,845 | 13,819,453 | 17,493,838 | 20,953,657 | 17,739,129 | 100,431,990 |
| **DLW CB4856 Hawaiian Chromosome Length (this study)** | 15,045,644 | 15,257,363 | 13,206,755 | 17,183,882 | 20,547,529 | 17,584,915 | 98,826,088 |
| **N2 Bristol Bases Aligned** | 15,100,574 | 15,303,320 | 13,222,676 | 17,330,119 | 20,947,147 | 17,738,394 | 99,642,230 (99.21%) |
| **% Syntenic Aligned Bases** | 93.31 | 88.56 | 90.61 | 95.42 | 87.04 | 98.73 | 92.23 |
| **SNPs\*** | 30,394 | 48,365 | 29,881 | 30,497 | 87,300 | 19,861 | 246,298 |
| **Indels\*** | 11,460 | 13,716 | 10,530 | 11,221 | 20,063 | 6,799 | 73,789 (275,442 bp) |
| **SVs** | 863 | 808 | 649 | 619 | 925 | 470 | 4,334 (2,654,902 bp) |
| **HDRs** | 185 | 270 | 165 | 138 | 356 | 60 | 1,174 (6,864,884 bp) |

\* All variants listed are only those for which the DLW CB4856 Hawaiian genome was homozygous

Analysis of universal single-copy orthologs (Simão et al. 2015; Manni et al. 2021) in our *de novo* N2 Bristol and CB4856 Hawaiian genomes revealed greater than 98% completeness (Supplemental Figure S1) and validate that our assemblies are high quality.

### *De novo* genome assemblies of the N2 Bristol and CB4856 Hawaiian isolates enhance detection of genomic variation

Previous comparisons of the genetic variation between N2 Bristol and CB4856 Hawaiian have relied on a short-read N2 Bristol reference genome (Thompson et al. 2015; Kim et al. 2019), and the amount of variation has yet to be re-assessed using a modern long-read N2 Bristol assembly. Utilizing our N2 Bristol and CB4856 Hawaiian strains, we aligned CB4856 Hawaiian short reads to our N2 Bristol assembly. This analysis revealed a total of 246,298 homozygous SNPs and 73,789 homozygous indels across the genome (Table 1, Figure 1C-D). While many of these SNPs and indels overlapped with gene annotations, they were under-enriched in gene sequences (Figure 1D-E, Supplemental Figure 2). To identify large sequence variants and chromosome rearrangements, we used whole-genome alignments (see Methods). We identified a total of 4,364 structural variants, which are categorized as insertions, deletions, and other chromosomal rearrangements spanning at least 50bp.

We also identified 1,174 Highly Divergent Regions (HDRs) (Goel et al. 2019) across the genome. HDRs are defined as regions of the genome over 50bp in length that result in low-quality pairwise alignments due to the presence of multiple gaps within these alignments (Goel et al. 2019). Overall, greater than 9.9% of the DLW N2 Bristol genome (∼10.0Mb) displayed variation through SNPs, indels, SVs, and HDRs when compared to the DLW CB4856 Hawaiian genome. SVs and HDRs represented only 1.3% and 0.3%, of variant sites between N2 Bristol and CB4856 Hawaiian respectively, but accounted for over 94% (9.5Mb) of sequence variation (Table1). Including heterozygous variants, our short-read analysis detected 3% more SNPs and 18% more indels than previously discovered using short-read assemblies of N2 Bristol and CB4856 Hawaiian (Thompson et al. 2015). Utilizing whole-genome alignment comparisons (Li 2018; Goel et al. 2019), we identified 985 more SV sites than previously reported (Nattestad and Schatz 2016; Kim et al. 2019). This increased sensitivity in variant site detection highlights the power of combining long-read and short-read sequencing to create accurate genome assemblies for comparative genomic studies.

Given an enhanced detection of variant sites between our N2 Bristol and CB4856 Hawaiian assemblies, we were interested in the genome-wide distribution of all variant sites. Given previous reports (Thompson et al. 2015; Kim et al. 2019), we expected a greater density of variation in the terminal thirds (the “arm-like” regions) of each chromosome. Indeed, there is a significant concentration of SNPs, indels, SVs and HDRs in the arm-like regions relative to the central region of each chromosome (Supplemental Figure S2). Over 78% of all SNPs, indels, SVs, and HDRs are in the arm-like domains of each chromosome (Genome-wide averages: 75.12% of SNPs, 78.24% of indels, 71.39% of SVs, 90.77% of HDRs). To determine if the enrichment of SNPs, indels, and SVs in the chromosomal arm-like regions was significant, we compared the observed distribution of each variant category with random permutations of each category of variant (Heger et al. 2013). SNPs, indels and HDRs on the autosomes were significantly enriched in the arm-like regions (SNPs: 1.36-1.77 fold enrichment; Indels: 1.47-1.84 fold enrichment; HDRs: 1.70-2.06 fold enrichment; p < 0.001 by hypergeometric test). SVs, however, were only significantly enriched on the arm-like regions of autosomes I, III, and IV (1.64-1.92 fold enrichment; p<.001 by hypergeometric test). The fold enrichment of all variants on the arm-like regions of the X chromosome was slightly weaker, ranging from 1.23-1.64 (SNPs: 1.26 fold enrichment; Indels: 1.26 fold enrichment; SVs: 1.23 fold enrichment; HDRs: 1.64 fold enrichment; all p-values < 0.05 by hypergeometric test). Similar to previous observations (Thompson et al. 2015), there were a few hyper-variable regions with a greater density of SNPs and short indels in the central regions of the autosomes, particularly on chromosomes IV and V (Figure 1 C-D).

Structural variations and HDRs account for most of the base-pairs affected by sequence divergence between our N2 Bristol and CB4856 Hawaiian lineages. The SVs identified ranged in size from 50bp to 592kb (Figure 2D-E), and HDRs ranged from 50bp to 199kb. Within the SVs detected, we identified 47 non-alignable structures, 2 duplications, 18 inversions, and 2 translocations. Non-alignable regions (NOTALs) are highly diverged regions containing many repeats and low-complexity sequences that are inhibitory to whole-genome alignment. From our whole-genome alignments of the DLW N2 Bristol and DLW CB4856 Hawaiian genomes, the non-alignable regions between the two genomes comprise 1.39Mb of sequence, ranged in size from 50-592kb, and comprise <0.5% of coding genes in the Bristol genome. One 156kb translocation was found on the right end of CB4856 Hawaiian chromosome V (V:15,871,614-16,027,614bp), while the other translocation, 38kb, was found to be inverted near a telomere of CB4856 Hawaiian chromosome IV (IV: 176:38,447bp). The largest duplication was found on Hawaiian chromosome III (III: 11,819,363-11,860,261). Together, our analyses provide improved variant site identification in wild isolate genomes and further illuminates previously undetected large structural variations and HDRs.

**Figure 2.**
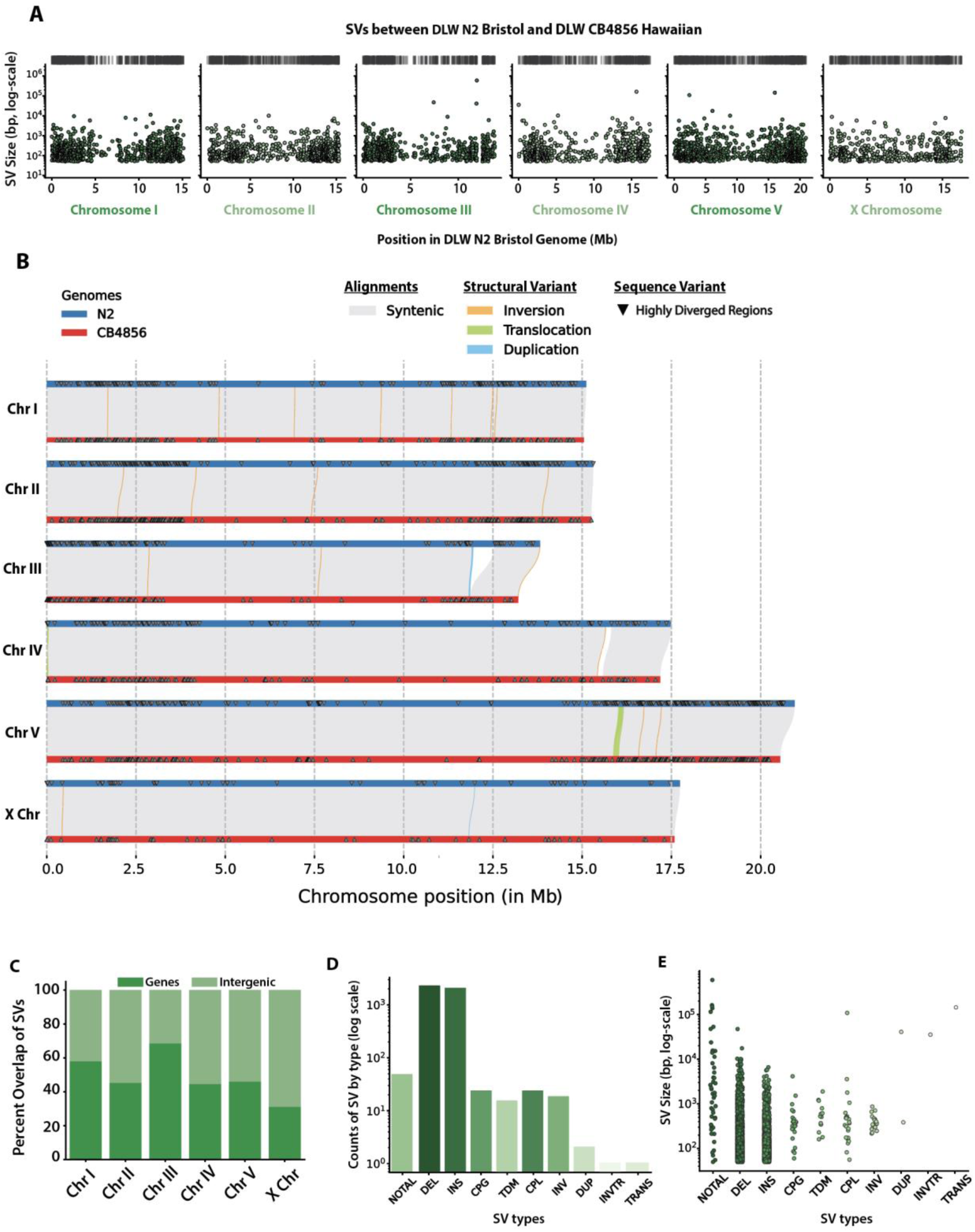
Genomic distribution and size of SVs between the DLW N2 Bristol and DLW CB4856 Hawaiian genomes. **(A)** Histograms depicting the distribution of SVs across each chromosome in 100kb bins. Black dashes above each histogram correspond to the genomic locations of SVs that are greater than 20kb in size. **(B)** Chromosome alignment plot depicting syntenic regions between N2 Bristol and CB4856 Hawaiian, structural variants, and highly divergent regions (HDRs). The width of lines showing SVs are proportional to their size. Only rearrangements 1kb or greater in size are shown. **(C)** Stacked bar plots showing the percentage of CB4856 Hawaiian SVs that overlap with intergenic and gene-coding regions of the DLW N2 Bristol genome. **(D)** Bar plots showing the number of each type of SV identified. **(E)** Strip plots showing the log-scaled size distribution of SVs separated by type. For SV types: NOTAL = non-aligned regions, DEL =deletion, INS = insertion, CPG = copy gain in query genome, CPL = copy loss in query genome, TDM = tandem repeat region, INV = inversion, DUP = duplication, TRANS = translocation, and INVTR = inverted translocation. For D and E, different colors only correspond to the different types of SV identified.

### SNPs, indels, and SVs are under-enriched in coding regions

Genes are enriched in the central region of all chromosomes in *C. elegans* (C. elegans Sequencing Consortium 1998), but there are some genes scattered across the chromosome arm-like regions. While much of the sequence variation is enriched in the arm-like regions, we wanted to know whether this variation was affecting coding sequences across the genome. Thus, we tested whether the SNPs, indels, SVs, and HDRs we identified between our N2 Bristol and CB4856 Hawaiian assemblies were enriched in genes versus intergenic space. Based on our remapped annotations (see Methods, LiftOff (Shumate and Salzberg 2021)), approximately 61.8% of the DLW N2 Bristol genome is comprised of gene sequences, with exons and introns representing 28.6% and 33.2% of the genome, respectively. Thus, we would expect corresponding proportions of each variant type to overlap within each annotation if variant sites were uniformly distributed across the genome. To determine if SNPs and indels were enriched in genes, we used the Genomic Association Tester (Heger et al. 2013) to compare the observed overlap of our variant sites in each remapped annotation to simulated uniform distributions of SNP and indel intervals. Fold enrichments represent the ratio of observed overlap to simulated overlaps, whereby a fold enrichment of 1.0 means there is no difference between the observed and simulated datasets. The greatest overlap of SNPs and indels in gene regions were observed on the autosomes (SNPs: 55.2-69.1%; indels: 52.1-68.5%), while only 49.1% of SNPs and 42.0% of indels were found in genes on the X chromosome (Figure 1 E-F). Across the genome, SNPs were slightly under-enriched in gene regions with an average fold enrichment of 0.96 (hypergeometric test, p-value <0.05). The average fold enrichment of indels in gene regions was lower than observed with SNPs (fold enrichment of 0.90, p-value <0.05), which could be due to selection against indels within coding regions. For SNPs and indels that did overlap with genes, intron sequences harbored the greatest amount of each variant type (SNPs: fold enrichment 1.14; indels: fold enrichment 1.45; Supplemental Figure 2). In conclusion, SNPs and indels are slightly overrepresented in intergenic regions of the *C. elegans* genome.

The distribution of SVs and HDRs across each chromosome resembles the genomic distribution of SNPs and indels (Figure 2A-B). To determine whether these large variant regions were enriched in intergenic versus coding regions, we compared the enrichment of simulated uniform distributions of SVs to those we identified. On the autosomes, 44.5-68.5% of SVs overlapped with gene regions compared 31.1% on the X chromosome (Figure 2C). Compared to SNPs and indels, structural variations on each chromosome, except chromosome III, displayed significant fold enrichments in intergenic regions (fold enrichments 1.3-1.5, p-values < 0.001; Supplementary Figure S2). Similar to SVs, highly divergent regions overlapped with 38.7-66.6% of genes on the autosomes and 24.3% on the X chromosome. HDRs were significantly enriched in intergenic regions of all chromosomes except chromosome I (fold enrichments 1.15-1.65; p-values < .05). Taken together, our data demonstrate that non-coding regions on the chromosome arm-like regions harbor most of the sequence variation between N2 Bristol and CB4856 Hawaiian lineages.

### Minimal movement of DNA transposons between the N2 Bristol and CB4856 Hawaiian lineages

Early analyses of the *C. elegans* genome indicated that approximately 12-16% of the genome is comprised of transposable elements (TEs) (*C. elegans* Sequencing Consortium 1998; Bessereau 2006), with *Tc1/mariner* elements as one of the most widely studied DNA transposons that can be active in laboratory strains (Emmons et al. 1983; Liao, Rosenzweig, and Hirsh 1983). While transposable element distributions have been assessed in wild *C. elegans* strains using older reference genomes and Illumina short-read sequencing (Laricchia et al. 2017), the complete TE composition has not yet been reassessed in a *de novo* assembly built from long-read sequencing. Further, new families of eukaryotic Class II transposons, have been discovered (Bao et al. 2009), and it remains unclear if these emerging families of DNA transposable elements comprise a significant proportion of the *C. elegans* genome.

To identify and locate known transposable element sequences in our N2 Bristol and CB4856 Hawaiian assembled genomes, we used a transposable element identification pipeline that applies an ensemble of programs to find all known RNA and DNA transposable element families (Riehl et al. 2022). We found that approximately 14.7 and 14.3% of our N2 Bristol and CB4856 Hawaiian assemblies, respectively, are composed of transposable element sequences (Supplemental Table 1). For both genome assemblies, the distribution of TEs was concentrated in the terminal third, arm-like regions of each chromosome (Figure 3A-B). Class II DNA TEs represented 96% of all TEs identified in each genome, and Zator elements are 52% of these Class II DNA TEs present in each genome (Supplemental Table 1, Figure 3C-D). To our knowledge, movement of Zator elements and other recently identified TE families has not yet been analyzed in *C. elegans* laboratory strains. Further, we also found that N2 Bristol genome has 20.6% more TEs from the *Tc1/mariner* family than the CB4856 Hawaiian genome (Supplemental Table 1).

**Figure 3.**
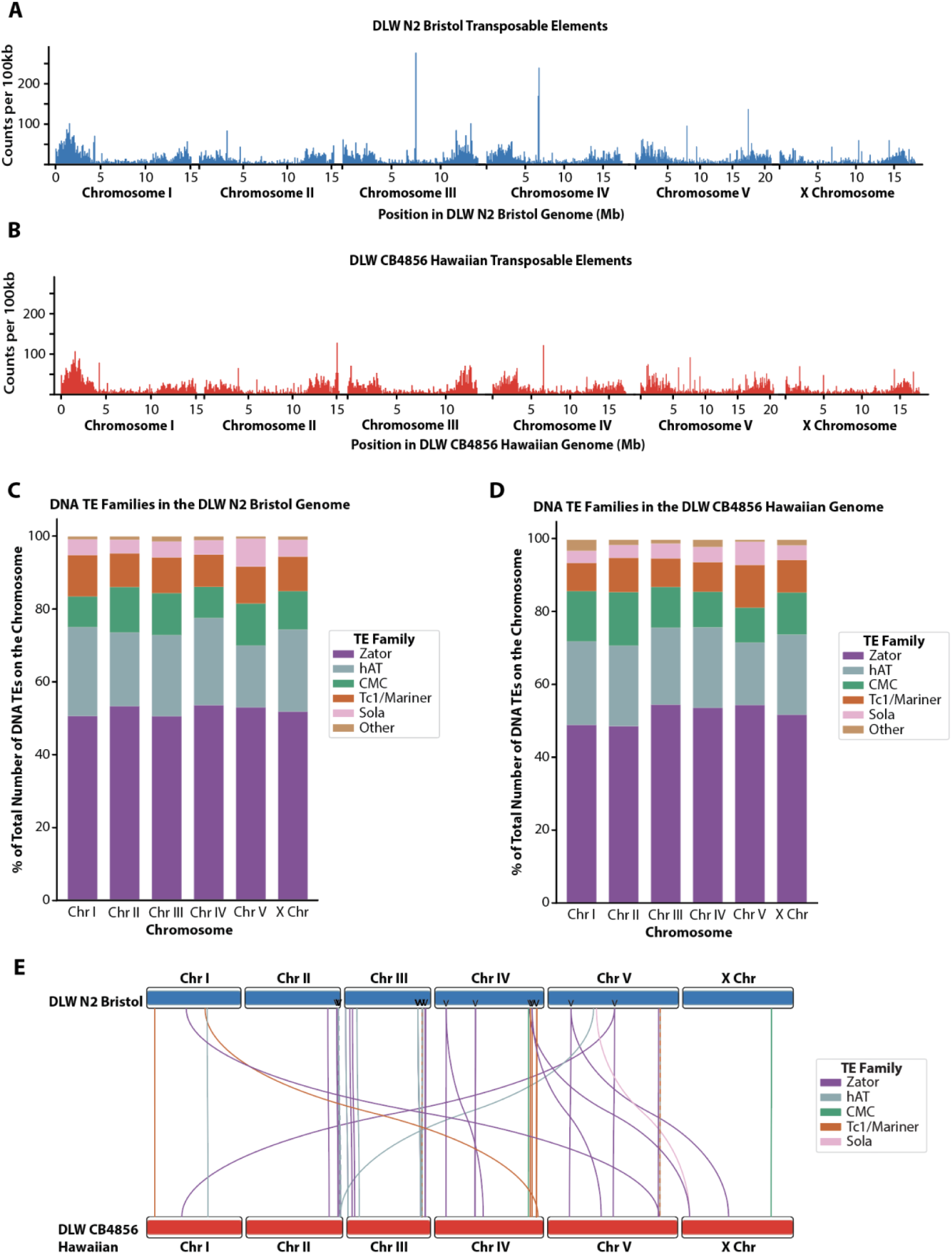
Genomic distributions of transposable elements in the DLW N2 Bristol and DLW CB4856 Hawaiian genomes. Histograms depicting the distributions of transposable elements across the DLW N2 Bristol genome in 100kb bins. **B**) Histograms depicting the distributions of transposable elements across the DLW CB4856 Hawaiian genome in 100kb bins. **C,D)** Stacked bar plot depicting the percent of total DNA transposable elements on DLW N2 Bristol (C) and DLW CB4856 Hawaiian (D) chromosomes accounted for by specific DNA transposon families. For TE families: CMC= CACTA, Mirage and Chapaev families; hAT = hobo and Activator families; Other = MITE, Novosib and Helitron families. **E**) Ideogram depicting the locations of individual DNA transposable elements that moved between the DLW N2 Bristol genome and the DLW CB4856 Hawaiian genome. DLW N2 Bristol chromosomes are represented by the blue boxes on the top, and DLW CB4856 Hawaiian chromosomes by the red boxes on the bottom. Each line represents an individual transposable element sequence, traced from its position on the DLW N2 Bristol genome to its unique position on the DLW CB4856 Hawaiian genome. Transposable elements predicted to have translocated are colored according to transposon class. Arrow heads across the Bristol N2 chromosomes indicate DNA TEs where duplicated copies are found in the Hawaiian CB4856 genome.

Since the N2 Bristol and CB4856 Hawaiian lineages were geographically isolated for thousands of generations, we sought to utilize our new TE annotation set to identify individual transposition events that occurred over the course of divergence between the two strains. Using whole-genome alignments and the SNPs we previously defined between these two lineages, we identified specific TE sequences with unique polymorphisms that enables individual transposons to be tracked between the N2 Bristol and CB4856 Hawaiian genome assemblies. Of the 18,392 total transposable elements identified in the N2 Bristol genome, 9,377 TEs were uniquely identifiable by sequence polymorphism. Among all N2 Bristol TEs with SNPs, only 1,535 elements were detectable in the CB4856 Hawaiian genome. While the vast majority of TEs were found to have not moved within either genome, we did identify 38 Class II DNA TEs that moved intrachromosomally and 9 TEs that moved interchromosomally (Figure 3E). Specifically, we detected 6 Zator elements and one each of *Tc1/mariner*, Sola, and hAT elements at different interchromosomal locations between the two lineages. In this analysis, we also found several unique copies of Class II DNA transposable elements in the N2 Bristol genome that had duplicated copies in the CB4856 Hawaiian genome (Figure 3E, arrowheads). While we were able to identify transposition events relative to the N2 Bristol genome, we cannot accurately infer the history of each CB4856 Hawaiian copy to determine which resulted from transposition versus duplication. Overall, the landscape of transposable elements remains largely unchanged across the history of divergence between the N2 Bristol and CB4856 Hawaiian lineages.

### Structural variants predominate the sequence divergence between lab strains

Much of the work exploring *C. elegans* genetic diversity utilizes comparisons of different natural isolates (Koch et al. 2000; Wicks et al. 2001; Thompson et al. 2015; Andersen et al. 2012). Work on germline mutation rates in *C. elegans*, however, suggest that considerable genetic variation may have been incurred during the laboratory setting (Denver et al. 2009). Given the rate of mutation accumulation in the germline (2.7 x 10^-9^ mutations per site per generation (Denver et al. 2009)) and a generation time of approximately three days, each N2 lineage alone may have accumulated up to ∼1,500 single nucleotide mutations since the 1970s, and nearly 790 potential mutations since the first genome was published in 1998 (C. elegans Sequencing Consortium 1998). Notably, this predicted variation does not include the accumulation of indels and structural variations. Thus, the N2 Bristol and CB4856 Hawaiian genomes present in each lab strain likely carries considerable genomic variation relative to other labs isolates. Previous studies using earlier genome assemblies identified many segmental duplications between lab lineages of wild type strains (Vergara et al. 2009). This variation may underpin phenotypic variation as well as previous work that has shown the lifespans of laboratory N2 Bristol isolates varies between 12-17 days (Gems and Riddle 2000). Taken together, accumulating evidence suggests that inter-lab genetic variation in wild type backgrounds may contribute to differences in experimental outcomes. High-quality lab-specific reference genomes may be an important tool to understand how genetics influences the phenotypes and processes studied by different laboratory groups.

To further evaluate the quality and differences of our genome assemblies, we aligned our N2 Bristol genome to the VC2010 Bristol (Yoshimura et al. 2019), as well as aligned our CB4856 Hawaiian genome to the Kim CB4856 Hawaiian genome (Kim et al. 2019). We expected that examining whole-genome alignments to previously validated long-read assemblies would reveal a striking degree of similarity. Comparing our N2 Bristol genome to VC2010, 99.9% of bases were alignable and 99.8% of bases were in syntenic alignments (Table 2). Analysis of our CB4856 Hawaiian genome versus the Kim CB4856 Hawaiian genome showed that 96.1% of bases were alignable, with 92.3% of bases in syntenic alignments (Table 3). This high degree of similarity within alignments gives us increased confidence in the quality of our own genome assemblies.

**Table 2.** Comparisons between the DLW N2 Bristol genome (this study) and VC2010 Bristol genome.

|  | Chromosomes |  |  |  |  |  | Total |
| --- | --- | --- | --- | --- | --- | --- | --- |
|  | I | II | III | IV | V | X |  |
| **DLW N2 Bristol Chromosome Length (this study)** | 15,114,068 | 15,311,845 | 13,819,453 | 17,493,838 | 20,953,657 | 17,739,129 | 100,431,990 bp |
| **VC2010 Bristol Chromosome Length (Yoshimura et al., 2019)** | 15,331,301 | 15,525,148 | 14,108,536 | 17,759,200 | 21,243,235 | 18,110,855 | 102,078,275 bp |
| **DLW N2 Bristol Bases Aligned** | 15,108,942 | 15,310,622 | 13,819,294 | 17,492,076 | 20,852,291 | 17,738,432 | 100,321,657 bp (99.89%) |
| **% Syntenic Aligned Bases** | 98.83 | 99.83 | 99.47 | 99.12 | 99.05 | 99.50 | 99.28 |
| **SNPs\*** | 169 | 124 | 164 | 209 | 280 | 216 | 1,162 |
| **Indels\*** | 150 | 261 | 210 | 262 | 378 | 267 | 1,528 (3465 bp) |
| **SVs** | 113 | 134 | 83 | 228 | 175 | 164 | 897 (2,010,282 bp) |
| **HDRs** | 8 | 14 | 10 | 21 | 24 | 11 | 88 (406,737 bp) |

\* All variants listed are only those for which the VC2010 Bristol genome was homozygous

**Table 3.** Comparisons between the DLW CB4856 Hawaiian genome (this study) and Kim CB4856 Hawaiian genome.

|  | Chromosomes |  |  |  |  |  | Total |
| --- | --- | --- | --- | --- | --- | --- | --- |
|  | I | II | III | IV | V | X |  |
| **DLW CB4856 Hawaiian Chromosome Length (this study)** | 15,045,644 | 15,257,363 | 13,206,755 | 17,183,882 | 20,547,529 | 17,584,915 | 98,826,088 |
| **Kim CB4856 Hawaiian Chromosome Length (Kim et al., 2019)** | 15,528,896 | 15,813,191 | 14,110,336 | 17,985,219 | 21,389,866 | 18,073,349 | 102,900,857 |
| **DLW CB4856 Hawaiian Bases Aligned** | 14,620,886 | 14,680,704 | 12,451,582 | 16,482,840 | 19,427,050 | 17,371,269 | 95,034,331 (96.16%) |
| **% Syntenic Aligned Bases** | 94.91 | 92.38 | 93.01 | 89.28 | 89.24 | 96.10 | 92.32 |
| **SNPs\*** | 60 | 52 | 71 | 108 | 135 | 115 | 541 |
| **Indels\*** | 240 | 190 | 175 | 242 | 238 | 213 | 1,298 (2157 bp) |
| **SVs** | 148 | 274 | 194 | 626 | 660 | 168 | 2,070 (6,923,335 bp) |
| **HDRs** | 19 | 70 | 25 | 100 | 144 | 12 | 370 (3,327,407 bp) |

\* All variants listed are only those for which the Kim CB4856 Hawaiian genome was homozygous

To assess how much genetic variation may exist between lab lineages of the most utilized wild-type strain, we first compared our N2 Bristol genome to VC2010 Bristol. We identified 1,162 homozygous SNPs and 1,528 homozygous indels. (Figure 4A-B, Table 2). In total, over 2.07Mb were affected by SNPs, indels and SVs, with 99.7% of this sequence divergence due to structural variations (Figure 4C, Table 2). While highly divergent regions have been observed between wild populations of *C. elegans* (D. Lee et al. 2021), we were also able to identify over 404kb of sequence as HDRs between these two laboratory Bristol lineages (Table 2). These HDRs identified between laboratory strains represent regions with multiple gaps between both genomes within a pairwise alignment in regions of synteny (Goel et al. 2019). In addition, we identified two inverted duplications (5.4kb and 12.9kb on chromosomes III and V, respectively) and 39 simple inversions. Four of these inversions are over 29kb in size and account for 11.6% of all structural variation between our N2 Bristol and the VC2010 Bristol genomes. SVs of this nature can be particularly disruptive to genome organization by impairing interactions between regulatory sequences or disrupting gene expression through loss of coding regions (Stranger et al. 2007; Hurles, Dermitzakis, and Tyler-Smith 2008).

**Figure 4.**
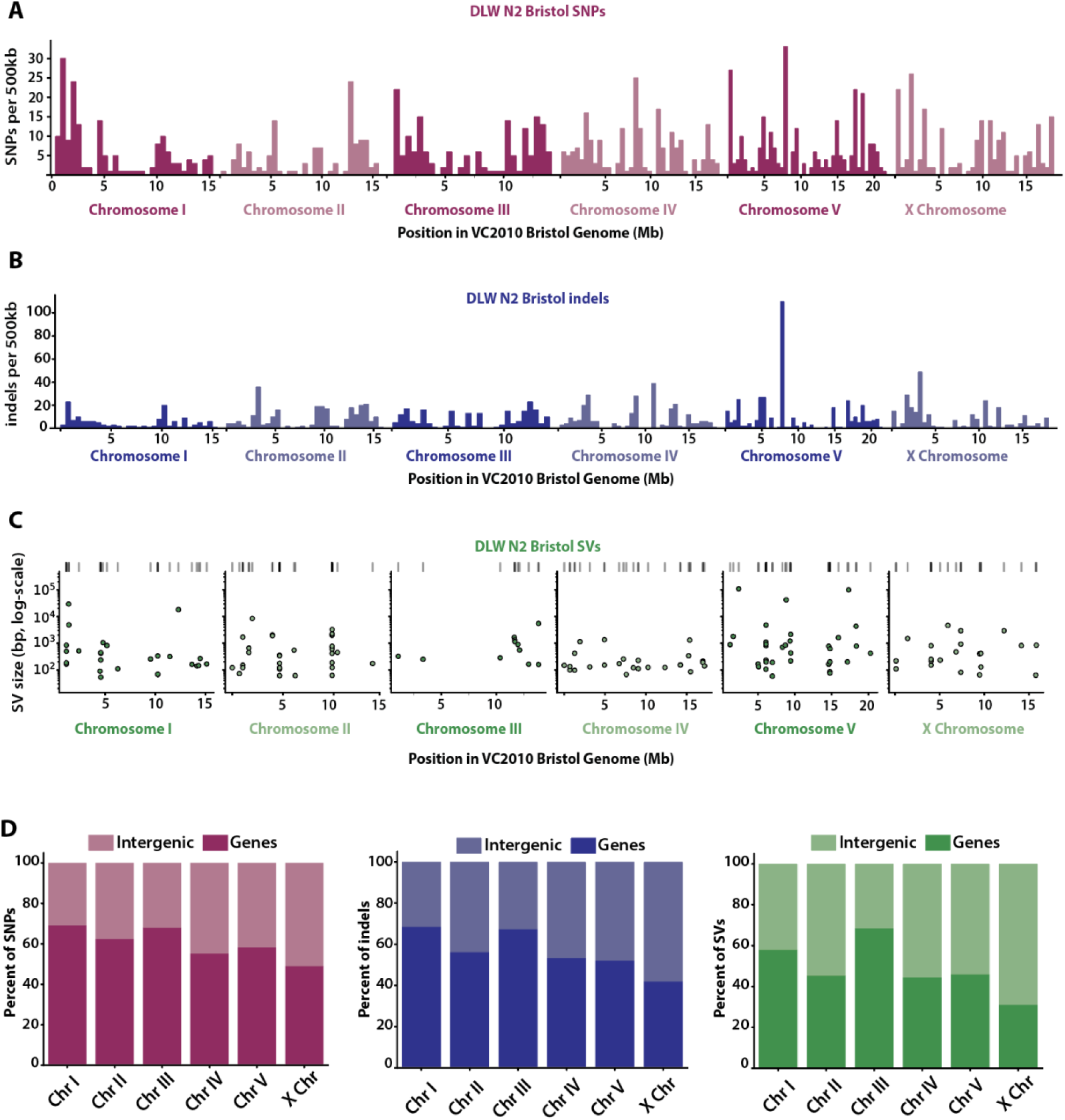
Genomic variation between the DLW N2 Bristol genome and the VC2010 Bristol genome. **(A-B)** Histograms depicting the distribution of DLW N2 Bristol SNPs and indels across each VC2010 Bristol chromosome in 500kb bins. **(C)** Scatterplots showing the genomic position of SVs with the log-scaled size of each SV on the y-axis. **(D)** The proportions of DLW N2 Bristol SNPs, indels, and SVs that overlap with intergenic versus gene-coding regions of the VC2010 Bristol genome.

Examination of our CB4856 Hawaiian lineage compared to the Kim *et al*., 2019 CB4856 Hawaiian assembly (Kim et al. 2019) revealed a greater amount of sequence divergence than comparisons between laboratory lineages of N2 Bristol. We identified 541 homozygous SNPs and 1,298 homozygous indels by aligning our CB4856 Hawaiian short reads to the Kim CB4856 Hawaiian genome (Supplementary Figure S3, Table 3, see Methods).

Notably, analysis of our whole-genome alignments identified over 9.5Mb of structural variation and HDRs between these two genomes. More than 66% of this structural variation, however, is due to unique, non-alignable regions. These non-alignable regions are highly divergent with many gaps in pairwise alignments that contain many repeats and low-complexity sequences. Further, over 3.3Mb in each Hawaiian genome falls within highly divergent regions. Taken together, we detected much more variation than anticipated between laboratory wild-type genomes. In the laboratory isolates of N2 Bristol and CB4856 Hawaiian, SVs affected the greatest number of base pairs, with a large portion of this variation due to large non-alignable regions, duplications, and inversions. Thus, the genomes of wild type strains present in some labs are not only unlike the most widely used reference genome in the *C. elegans* research community, but there are likely many large inter-lab genomic variations that might underlie some of the phenotypic differences observed in laboratory strains.

### Intergenic enrichment of variant sites between lab lineages of N2 Bristol and CB4856 Hawaiian

To determine whether specific genomic regions of lab strains are susceptible to sequence variation, we repeated our analysis of the genomic distributions of each variant class and assessed the enrichment of variant sites in gene annotations. We used LiftOff to remap the pre-existing Bristol gene annotations onto both the VC2010 Bristol and Kim CB4856 Hawaiian assemblies with similar success. To address whether the sequence divergence is nonrandomly enriched in each region of interest, we again used GAT to simulate random SNP, indel, SV, and HDR intervals 20,000 times and compared the simulated overlap to what we observed in between genomes. Notably, the stereotypical “arms”-vs-“center” genomic distribution of variants seen when comparing Bristol and Hawaiian genomes is not true for all chromosomes when comparing our Bristol genome to VC2010 (Figure 4A-C), with some chromosomes displaying a concentration of variation in the central region. SNPs were 1.2-1.5 fold enriched in the arm-like regions of chromosomes I, II, III, and V (p-values <0.01). Indels, however, were concentrated in the arm-like regions of all chromosomes with fold enrichments ranging from 1.2-1.5 (p-values < 0.05). While SVs were 1.2-2.2 fold enriched in the arm-like regions of each chromosome, this enrichment was only significantly higher than expected by null distributions on chromosomes I, IV, and the X chromosome (Supplemental Figure S4). Further, HDRs between Bristol lineages were 1.6 fold enriched on the arm-like regions of chromosome II and the X chromosome (p-values < 0.05), while displaying significant 1.8-2.1 fold enrichments in the center regions of chromosomes I, IV, and V (p-values < 0.01). Finally, we also wanted to determine whether the variant sites we detected between lab strains impacted gene coding regions. Between the two Bristol lineages, SNPs, indels, SVs, and HDRs were all under-enriched in gene coding regions and displayed significant enrichments in intergenic regions on most chromosomes (Supplemental Figure S4). Thus, variation between laboratory Bristol lineages is largely concentrated in non-coding regions of each chromosome.

We next examined the genomic distribution and enrichment of variant sites in gene annotations of the two Hawaiian genomes to see if the patterns of enrichment were similar to the Bristol genomes. After examining the enrichment of all variant types in both the arm-like regions and “centers” of each chromosome, it was clear that most chromosomes were enriched for variant sites in the arm-like regions and intergenic sequences, with a few exceptions as follows. (Supplemental Figure S5). SNPs were only enriched 0.84 fold in the arm-like regions of the X chromosome, and indels were enriched 0.81 fold in the arm-like regions of chromosome IV (p-values < 0.05). SVs were 0.64 fold enriched in the arm-like regions of chromosome II, and HDRs were 0.12 fold enriched in the arm-like regions of chromosome I (p-values < 0.001). We then examined the enrichment of all variants in intergenic versus gene sequences between the two CB4856 genomes. SNPs and indels showed significant 1.7-2.8 fold enrichments in intergenic regions on all chromosomes (p-values < 0.001). SVs displayed a significant 1.2-1.7 fold enrichment in the intergenic regions of all chromosomes. HDRs were 1.2-1.7 fold enriched in the intergenic regions of chromosomes II, III, IV, V and the X chromosome (all p-values < 0.05). In conclusion, analysis of the genetic variation between respective lab lineages of N2 Bristol and CB4856 Hawaiian revealed a striking amount of variation often present in intergenic sequences, with some weak enrichments in the arm-like regions versus the central regions of chromosomes.

## Discussion

Detection and characterization of sequence variation between individuals or across species is fundamental to our functional understanding of genomic elements and consequences of variation. Since the first draft of the *C. elegans* genome was released in 1998, the highly divergent strains N2 Bristol and CB4856 Hawaiian have been used extensively for comparative genomics studies(C. elegans Sequencing Consortium 1998; Koch et al. 2000; Wicks et al. 2001; Maydan et al. 2010; Andersen et al. 2012; D. Lee et al. 2021). The combined usage of short and long read sequencing to assemble genomes and to compare them has both increased the quality of our reference genomes as well as enhanced the genome-wide detection of sequence variants, new genes, and new genomic regions (Yoshimura et al. 2019; C. Kim et al. 2019; B. Y. Kim et al. 2021; Sarsani et al. 2019). In this study, we generate *de novo* assemblies for the N2 Bristol and CB4856 Hawaiian *C. elegans* isolates from our lab lineage using short-read and long-read sequencing. Our examination of the inter-lab genetic drift among wild-type strains suggests genomic analyses can be improved by resequencing the genomes of labs’ wild-type strains or utilizing strains with recently published, accurate genome assemblies. This also presents a strong argument for labs utilizing *C. elegans* in their research to frequently return to cryogenically preserved stocks of their wild type strains. These genomes will serve as additional tools for future comparative genomics studies, especially in the functional characterization of structural variations identified through whole-genome alignments.

### Genome assembly and genomic divergence in laboratory isolates

Earlier studies uncovering phenotypic and genetic variations between lab wild-type strains indicated that there are likely many underlying large-scale genomic differences (Denver et al. 2009; Vergara et al. 2009; Gems and Riddle 2000). Here we identify numerous SNPs, indels, SVs, and HDRs between different lab lineages of each wild isolate. The total amount of genomic variation is at levels higher than predicted by earlier mutation accumulation studies. Much of this variation, however, is due to SVs and HDRs, which have only recently become a detailed subject of study (Thompson et al. 2015; Kim et al. 2019; Lee et al. 2021). Our genome assemblies of the Bristol and Hawaiian strains corroborate prior results indicating that genomic variation is enriched in the distal arm-like regions of chromosomes between these natural isolates. Evolutionary genomic analysis has shown that recombination in the arm-like regions of each chromosome and balancing selection likely have shaped this landscape of sequence divergence across the 30,000-50,000 generations these strains have been geographically isolated (Thomas et al. 2015; Kern and Hahn 2018). In contrast, we find that the distribution of variant sites across the arm-like regions versus center domains of each chromosome between lab lineages is not as strong or consistent as seen when comparing N2 Bristol to CB4856 Hawaiian genomes. This result could indicate that in relatively short timescales (∼3,000-5,800 generations), selection for the accumulation of mutations in the arm-like regions, particularly in noncoding regions, is not sufficient to consistently eliminate sequence divergence away from the gene-dense chromosome centers. Further, we found that SNPs, indels, and structural variations were highly enriched in intergenic regions when comparing the genomes of laboratory strains. Although many of the sequence variants we identified are not directly disrupting coding sequences, it remains possible that genetic drift in these regions are altering the function of intergenic regulatory sequences such as promoters and enhancers. Thus, the accumulation of disruptive genomic changes within regulatory regions in the gene-dense centers of chromosomes may underpin many of the phenotypic differences observed in laboratory wild-type strains, such as variance in lifespan (Gems and Riddle 2000).

### Highly variable arm-like domains on *C. elegans* chromosomes

The arm-like regions of *C. elegans* chromosomes exhibit a striking degree of variation that is highly correlated with large domains of increased recombination, which is a pattern observed in many species (Andersen et al. 2012; D. Lee et al. 2021; Kern and Hahn 2018; Rockman and Kruglyak 2009). In *C. elegans,* these divergent autosomal arm-like domains coincide with a disproportionate fraction of newer, rapidly evolving genes as compared to the center regions of each chromosome, which house highly conserved essential genes (C. elegans Sequencing Consortium 1998; Kamath et al. 2003). The development of new tools to detect larger structural variations through alignment of assemblies or long sequencing reads has revealed many SVs on the chromosomal arm-like domains (Mahmoud et al. 2019; C. Kim et al. 2019). The fact that SVs are enriched in the arm-like regions, which also display elevated levels of recombination, is notable given the fact that large structural variants such as inversion are typically inhibitory to recombination (Miller, Cook, and Hawley 2019). The arm-like regions of *C. elegans* chromosomes are enriched for many repetitive elements, including transposable elements, tandem repeats, and low complexity repeat sequences (C. elegans Sequencing Consortium 1998; Surzycki and Belknap 2000). The presence of many SVs in the arm-like regions could be due to errors in double-strand DNA break repair and heterologous recombination in regions adjacent to highly repetitive sequences, thereby causing chromosomal rearrangements. Similar rearrangement events are known to contribute to many human genomic disorders like Prader-Willy Syndrome or Charcot-Marie-Tooth disease (Carvalho and Lupski 2016; Stankiewicz and Lupski 2010). Future investigations assessing the occurrence of SVs adjacent to highly repetitive regions and sites of homologous recombination will be invaluable in understanding how differences in genomic organization arise between divergent lineages of *C. elegans*.

With regard to genomic rearrangements and their impact on genome function, renewed attention must be given to the contribution of transposable elements and their mobility within and between chromosomes. While Sola and Zator elements are relatively recent in their discovery within *C. elegans* and other eukaryotic genomes (Bao et al. 2009; Riehl et al. 2022), our data suggests there may be many active TE copies in these families, particularly Zator elements. Historically, much attention has been given to the impact of *Tc1/Mariner* transposition on genomic architecture, but the contribution of Zator elements to changes in genome structure and gene regulation merits further future investigation. Our analysis of TE mobility only examines two endpoints across the long period of divergence between the Bristol and Hawaiian lineages. It remains unclear, however, whether many of these newly characterized TEs remain active and whether they contribute to the growing catalog phenotypic differences displayed between laboratory lineages of Bristol and Hawaiian *C. elegans*.

Finally, the generation of multiple independent long read *de novo* genome assemblies for both N2 Bristol and CB4856 Hawaiian isolates provides a powerful toolkit for comparative genomics and evolution studies. Many prior studies assessing the *C. elegans* recombination landscape have relied on mapping recombination in worms heterozygous for Bristol and Hawaiian chromosomes. The high sequence divergence and large structural variations between Bristol and Hawaiian which we describe, however, may have positional impact on the distributions of crossover sites. Our identification of variants in Bristol strains enables polymorphism mapping by crossing different lab-lineages of N2 Bristol, avoiding the potential confounding effects of crosses with other wild isolates. Additionally, further identification and functional characterization of polymorphic sites and structural variations present between lab lineages of N2 Bristol and CB4856 Hawaiian could provide new insights into how pronounced phenotypic differences in the lifespan, feeding behavior, and reproductive fitness arise in modern lab-derived strains (Gems and Riddle 2000; Zhao et al. 2018). To summarize, we demonstrate the importance of using long and short-read sequencing to generate modern reference genome assemblies and maximally detect sequence variation, while highlighting the potential genomic underpinnings of phenotypic variations in laboratory lineages of *C. elegans*.

## Methods

### *C. elegans* culture and sucrose floatation

The N2 Bristol and CB4856 Hawaiian strains of *C. elegans* were grown at 20°C on standard NGM agar plates seeded with the OP50 strain of *E. coli* as a food source. To minimize bacterial contamination in downstream gDNA sample preps, we performed sucrose floatation on pooled populations of each isolate. Worms were washed from plates with 8mL cold M9 buffer and transferred to 15mL glass centrifuge tubes using a glass Pasteur pipette. Collected worms were centrifuged at 3000rpm at 4°C and washed in 4mL of fresh M9 twice. To separate worms from bacteria and other debris, 4mL of 60% sucrose solution was added to 4mL of M9 buffer and worms and vortexed briefly. The mixture was then spun at 5000 rpm at 4°C for 5 minutes. Using a glass pipette, the floating layer of worms were transferred to a new glass centrifuge tube on ice and brought up to 4mL in fresh M9. Worms were then incubated at room temp for 30 minutes and gently vortexed every 5 minutes. Worms were washed three times in equal volume of fresh M9 were performed before storing collected worms in M9 at 20°C before genomic DNA (gDNA) extraction.

### Long-read and short-read sequencing

Genomic DNA was extracted from worms using the Qiagen DNeasy Blood and Tissue Kit. Sequencing was performed on pooled populations of N2 and CB4856 after reducing bacterial contamination by sucrose float for each strain. For PacBio long-read sequencing, library preparation was performed on pooled populations of worms for each isolate by the University of Oregon’s Genomics and Cell Characterization Core Facility and sequenced on the Sequel II system. For Illumina short-read sequencing, library preparation was performed on pooled populations of worms for each isolate by the University of Oregon’s Genomics and Cell Characterization Core Facility. The short-read libraries were then sequenced on an Illumina HiSeq4000 (2 x 150bp).

### Long-read genome assembly and short-read refinement

PacBio long-reads were aligned to the E. coli genome using BWA (Li and Durbin 2009) (version 0.7.17), and reads that aligned to the bacterial genome were removed. De novo genome assembly was performed for N2 Bristol and CB4856 Hawaiian using Canu (Koren et al. 2017) (version 1.7). To refine the long-read assemblies, short-reads from each isolate were aligned to their respective long-read assembly using BWA-MEM (version 0.7.17). Aligned reads in SAM format were sorted and converted to BAM format using SAMtools(Li et al. 2009). Using Picard (https://broadinstitute.github.io/picard/), read groups were added via AddOrReplaceReadGroups, and duplicate reads were filtered using MarkDuplicates. Some bases may have been inaccurately called due to lower sequencing coverage, larger error rate in PacBio sequencing, or predominating alleles present in the population of each isolate that could be revealed by greater sequencing depth afforded by Illumina sequencing. GATK’s HaplotypeCaller (McKenna et al. 2010) and Freebayes (Garrison and Marth 2012) were utilized to generate VCF files representing potentially inaccurate sites in each initial assembly. Coverage thresholds were manually determined using IGV for each assembly. Sites were filtered according to manual values using VCFtools (Danecek et al. 2011; Danecek and McCarthy 2017)). Error correction was performed on single-nucleotide alleles using BCFtools *consensus* (Danecek and McCarthy 2017) and alternate indel alleles. After filtering potential sites by sequencing depth thresholds determined for each chromosome, this left 4237 and 36145 corrections for the N2 Bristol and CB4856 Hawaiian genomes, respectively. Of these sites, less than .7% were unable to be resolved, and all of these were short indels comprising less than .001% of each genome.

### Assessing genome assembly completeness

To further assess the quality and completeness of our N2 Bristol and CB4856 Hawaiian assemblies, we used BUSCO (Simão et al. 2015; Manni et al. 2021). BUSCO was run in a Docker container (https://busco.ezlab.org/busco_userguide.html) in genome mode. For each assembly, the quality and presence of expected orthologous genes was checked against the nematoda and metazoan lineage databases.

### SNP and indel Calling in N2 and CB4856 assemblies

Illumina short reads from the DLW N2 Bristol and DLW CB4856 Hawaiian genome were trimmed using Trimmomatic (Bolger, Lohse, and Usadel 2014) to remove adapter and barcode sequences. The trimmed CB4856 reads were then aligned to the DLW N2 Bristol reference genome using BWA-MEM so that SNPs and indels present between N2 Bristol and CB4856 Hawaiian could be identified. All resulting variant positions comparing our N2 Bristol and CB4856 Hawaiian genomes are in relation to the N2 Bristol assembly. Aligned reads in SAM format were then sorted using SAMtools (Li et al. 2009) and converted to BAM files. Using Picard read groups were added via AddOrReplaceReadGroups, and duplicate reads were filtered using MarkDuplicates as described above. BAM files with filtered duplicate reads were used to call variants using a combination of GATK HaplotypeCaller, Freebayes, and BCFtools. The three resulting VCF files containing SNPs and indels were then concatenated, further filtered for duplicate sites and low-quality variants, and sorted using BCFtools. SNPs with QUAL scores of 30 or greater, a minimum of 10 variant reads, and a minimum of 30 total, high-quality reads were retained. To draw comparisons between other N2 assemblies, the most recent gene annotations were downloaded from WormBase (*C. elegans* VC2010, PRJEB28388). For comparisons between CB4856 Hawaiian genomes, assemblies were downloaded WormBase (*C. elegans* CB4856, PRJNA275000) and the NCBI BioProject Database under accession number PRJNA523481. To call variants between our N2 Bristol and CB4856 Hawaiian assemblies and those generated by other labs, short reads were aligned to the respective genomes and the SNP and indel calling pipeline was repeated as described above.

### Calling Structural Variants using whole-genome alignments

All assembly-to-assembly alignments were performed using Minimap2 (Li 2018). SyRI (Goel et al. 2019) was then used to parse the resulting SAM files and call structural variants and highly divergent regions (Structural rearrangements were plotted with the aid of Plotsr within the SyRI package. “NOTAL” or non-alignable regions in each genome were retained as SVs. To acquire NOTAL regions in each query genome, the Minimap2 alignment was repeated with the original reference and query genomes swapped. The sizes of HDRs depicted in Tables 1-3 are sizes relative to the reference genome in each comparison (*i.e*. N2 Bristol in Table 1). When comparing our CB4856 Hawaiian genome to the Kim CB4856 genome, 89% of the size difference in assemblies can be accounted for in the net sequence gained from Kim HDRs and unique NOTAL structures. NOTAL structures and gap-adjacent sequences in the Kim CB4856 genome are 1.5 and 1.6-fold enriched for low complexity and repeat sequences, respectively. These regions and sequence features are challenging for genome assembly and likely explain megabase-scale differences in genome assembly sizes.

### Converting gene annotations between assemblies

We converted gene annotations from the N2 reference assembly (cel235) to our N2 Bristol and CB4856 Hawaiian assemblies, as well as the VC2010 Bristol and Kim CB4856 Hawaiian assemblies. The gene annotations for the WBcel235 genome assembly were downloaded in GFF3 format from Ensembl (http://ftp.ensembl.org/pub/release-105/gff3/caenorhabditis_elegans/). Unlike previously established tools that require pre-generated chain files(James et al. 2003), Liftoff (Shumate and Salzberg 2021) can accurately remap gene annotations onto newly generated assemblies using Minimap2 assembly-to-assembly alignments. Rather than aligning whole genomes, Liftoff aligns only regions listed in the annotation files so that genes may be remapped even if there are large structural variations between two genomes. The Liftoff program was then used to remap annotations between the WBcel235 assembly onto each new genome assembly for N2 Bristol and CB4856 Hawaiian).

### Testing the association between variant sites and gene annotations

For each chromosome, to determine whether SNPs or indels were enriched within gene annotations, fold enrichment analyses were performed using the genomic association tester (GAT) (Heger et al. 2013) tool (https://github.com/AndreasHeger/gat.git). The observed enrichment of each variant type in gene annotations was compared to overlaps in simulated distributions SNPs or indels. Simulated distributions were created using 20,000 iterations whereby each variant type was randomly and uniformly distributed across each chromosome. SNPs and indel distributions were compared against intergenic, gene, intron, exon, and UTR annotations. Comparing the observed enrichment to the simulated distributions, statistical significance was assigned to the observed fold enrichment with p-values calculated from a hypergeometric test calculated within GAT. Per-chromosome BED files for SNP intervals were created from their original VCF using AWK. Per-chromosome BED files for indel intervals were calculated using a custom script. The GFF3 formatted annotations generated via liftoff were then broken down by chromosome, gene, exon, and UTR regions. Because intron regions were not explicitly written into each GFF3 file, they were calculated using BEDtools (Quinlan and Hall 2010). First, a joint BED file containing the UTR and exon regions were made using awk and sorted first by chromosome then by position. Using BEDtools these intervals were combined, and intronic regions were calculated by finding regions in gene intervals not covered by either UTR or exons. Intergenic spaces on each chromosome were calculated with the gene BED files and chromosome sizes as inputs. GAT was then run for each chromosome in each assembly.

### Transposable Element Identification and Tracking

The TransposonUltimate pipeline (Riehl et al. 2022) was run for both our N2 Bristol and CB4856 Hawaiian genome assemblies. MUST and SINE finder were run independently and integrated into the filtering steps of the pipeline manually. Additionally, we added LTR retriever to the TE identification ensemble to supplement LTR harvest and LTR finder. TE sequences that overlapped with SNPs were identified using BEDtools. CB4856 Hawaiian SNPs were applied to corresponding N2 Bristol TE sequences, and these sequences were cross-referenced with the original TransposonUltimate output for CB4856 Hawaiian for matches. Unique polymorphic TE sequences found in both genomes were then assessed for translocation events by examining genomic start coordinates in each genome. Utilizing whole-genome alignments for each chromosome, TEs were predicted to have moved if starting coordinates for each TE pair were did not correspond to relative changes in coordinates due to alignment.

## Data Availability Statement

The PacBio long-read and the Illumina short-read data generated in this study have been submitted to the NCBI BioProject database (https://www.ncbi.nlm.nih.gov/bioproject/) under accession number PRJNA907379. All custom scripts are available upon request. Strains are available upon request.

## Acknowledgements

We thank N. Kurhanewicz, C. Cahoon, E. Toraason, and J. Conery in the Libuda Lab for thoughtful discussion and comments on the manuscript. We are grateful to the University of Oregon’s Genomics and Cell Characterization Core Facility for sample prep and sequencing of our N2 Bristol and CB4856 Hawaiian strains. Over the course of many years beyond the time of this study, thanks for inspiration from Walter Rogers and invaluable guidance from Dr. Glenn C. Rowe given to Z.D.B.

## Funding

This work was supported by the National Institutes of Health T32HD007348 to Z.D.B and National Institutes of Health R35GM128890 and University of Oregon start-up funds to D.E.L.

## Conflict of Interest

The authors declare no conflicts of interest.

## Author Contributions

Z.D.B. polished primary genome assemblies, assessed genome quality, annotated the genomes, and analyzed genomic variation and structures. A.F.S.N. helped conceive this study and developed protocols for DNA purification, short-read Illumina sequencing, and variant calling. D.D. developed protocols for long-read sequencing, devised strategies for genome assembly, assembled the contigs and primary assembly, and filled gaps. C.A. identified transposable elements and tracked copies containing SNPs between genome assemblies. K.J.H. helped conceive this study, develop protocols for DNA purification, and purified the DNA for the long-read PacBio sequencing. D.E.L. helped conceive this study, led discussions for the comparison of different genomes, and coordinated work. All authors contributed to the manuscript, with most of the writing by Z.D.B., A.S.F.N., and D.E.L.

---

**Supplemental Figure S1.**
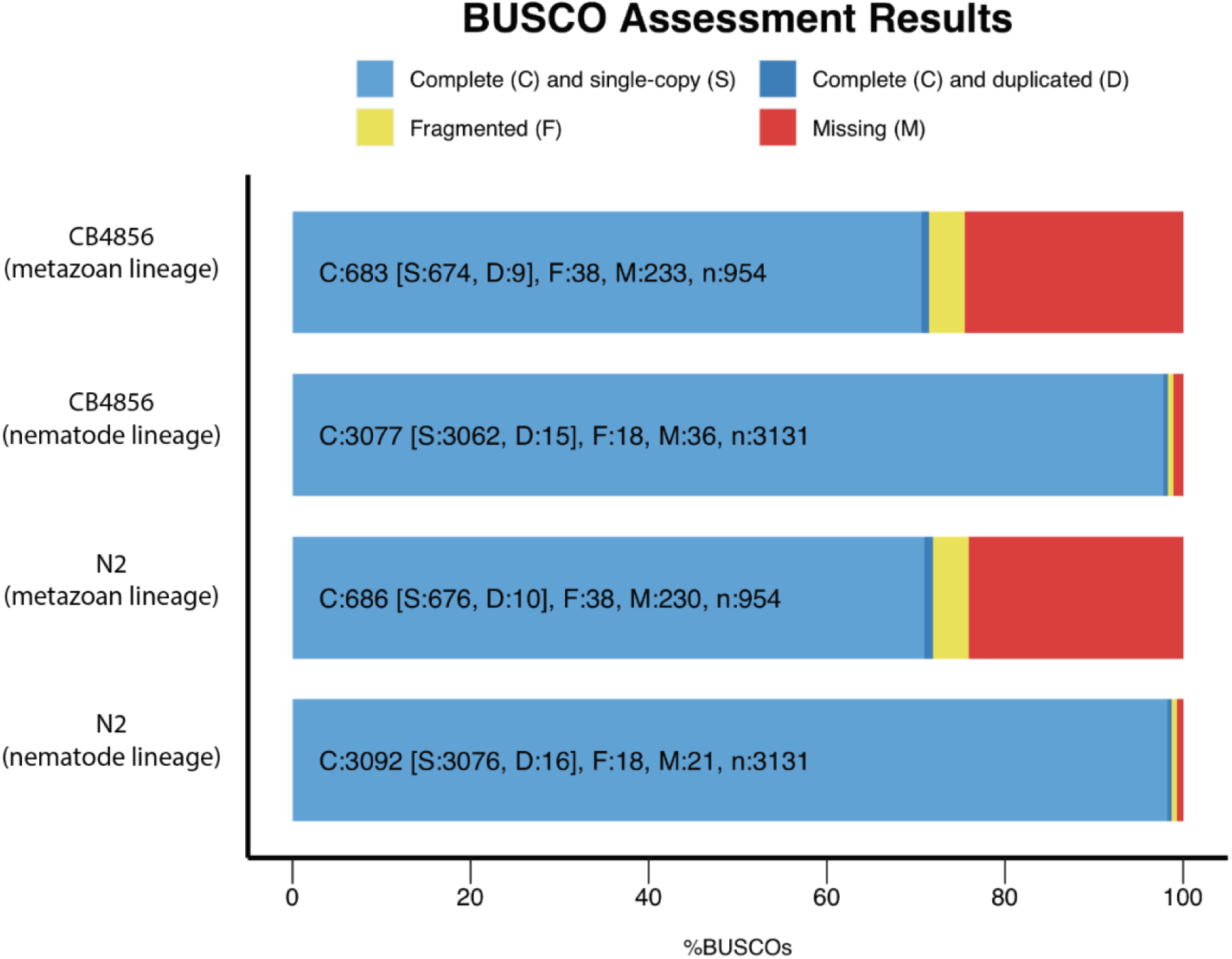
BUSCO analysis of the DLW N2 Bristol and DLW CB4856 Hawaiian genome assemblies. The presence of orthologous genes from metazoan and nematode lineages are shown for each genome assembly. Each orthologous gene analyzed is depicted as either Complete (C, blues), Fragmented (F, yellow), or Missing (M, red). Complete orthologs are then further categorized as single-copy (S, light blue) or duplicated (D, dark blue).

**Supplemental Figure S2.**
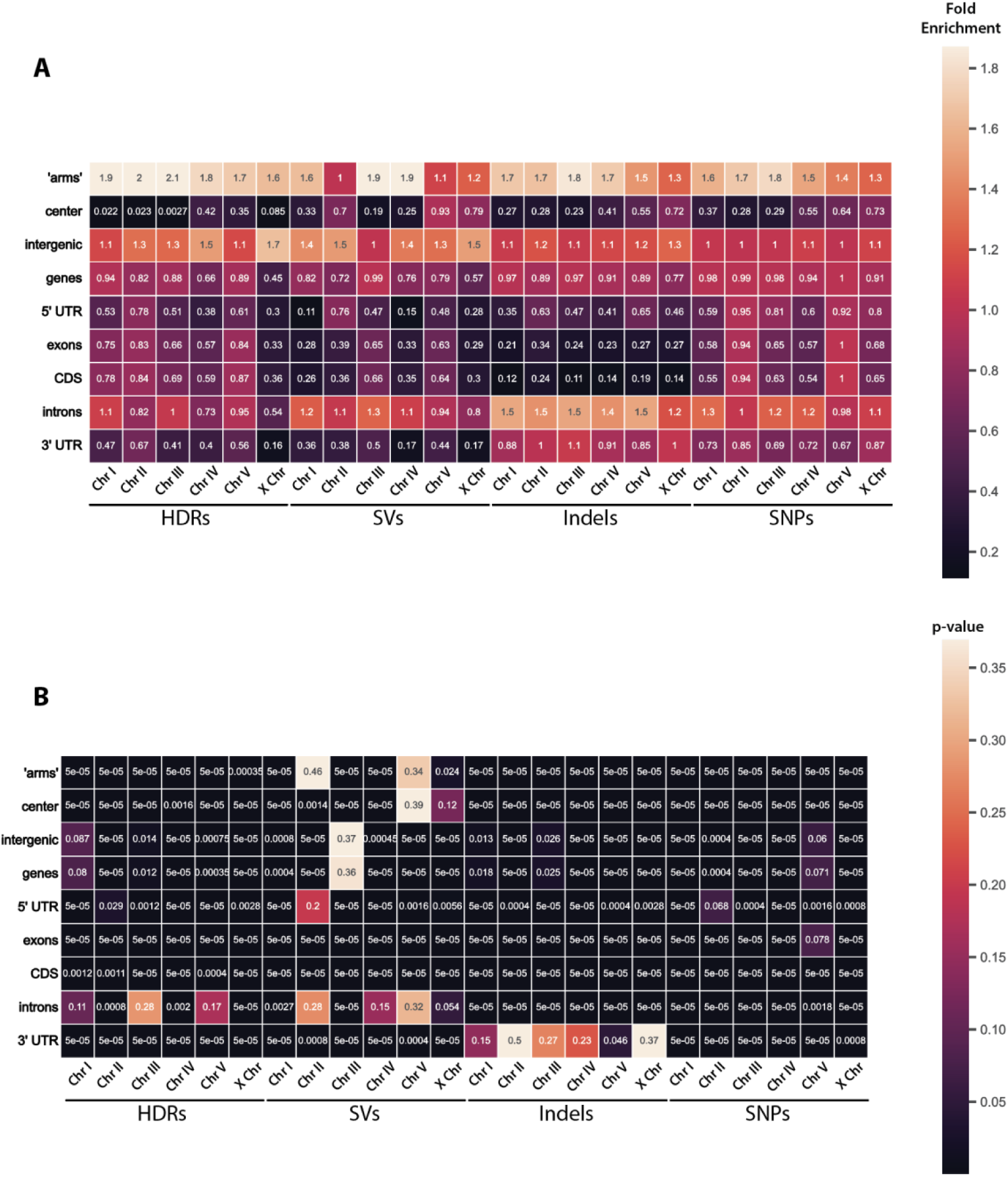
GAT interval-association test results analyzing the overlap of DLW CB4856 Hawaiian SNPs, indels, and SVs with N2 genome annotations. A) Heatmap showing the fold enrichment of each variant type within gene annotations for each chromosome. B) Heatmap of p-values associated with corresponding fold enrichments shown in panel A calculated by the hypergeometric test.

**Supplementary Figure S3.**
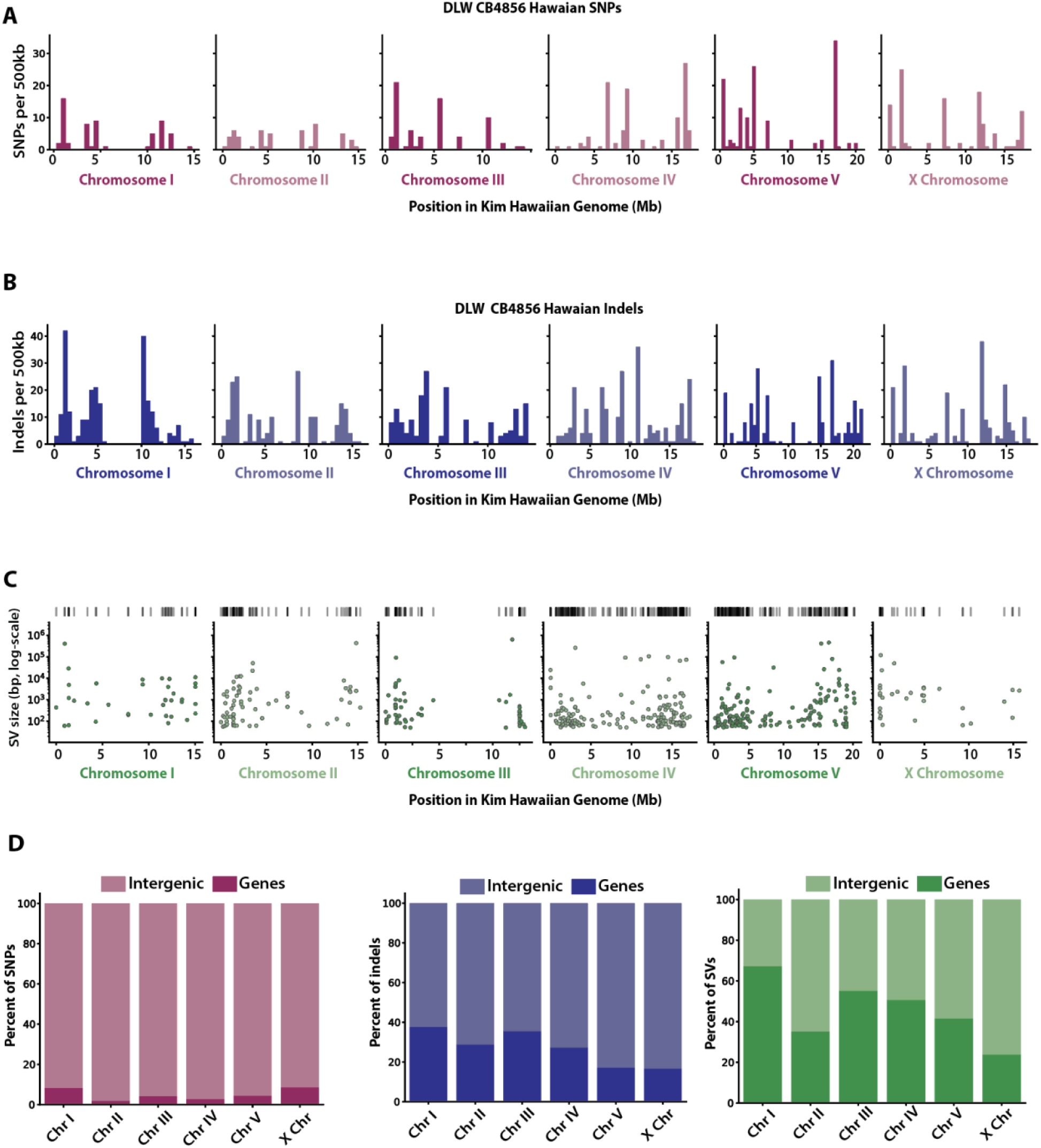
Genomic variation between the DLW CB4856 Hawaiian genome and the Kim CB4856 Hawaiian genome. (A-B) Histograms depicting the distribution of SNPs and indels across each Kim CB4856 Hawaiian chromosome in 500kb bins. (C) Scatterplots showing the genomic position of SVs with the log-scaled size of each SV on the y-axis. (D) The proportions of DLW CB4856 Hawaiian SNPs, indels, and SVs that overlap with intergenic versus gene-coding regions of the Kim CB4856 Hawaiian genome.

**Supplemental Figure S4.**
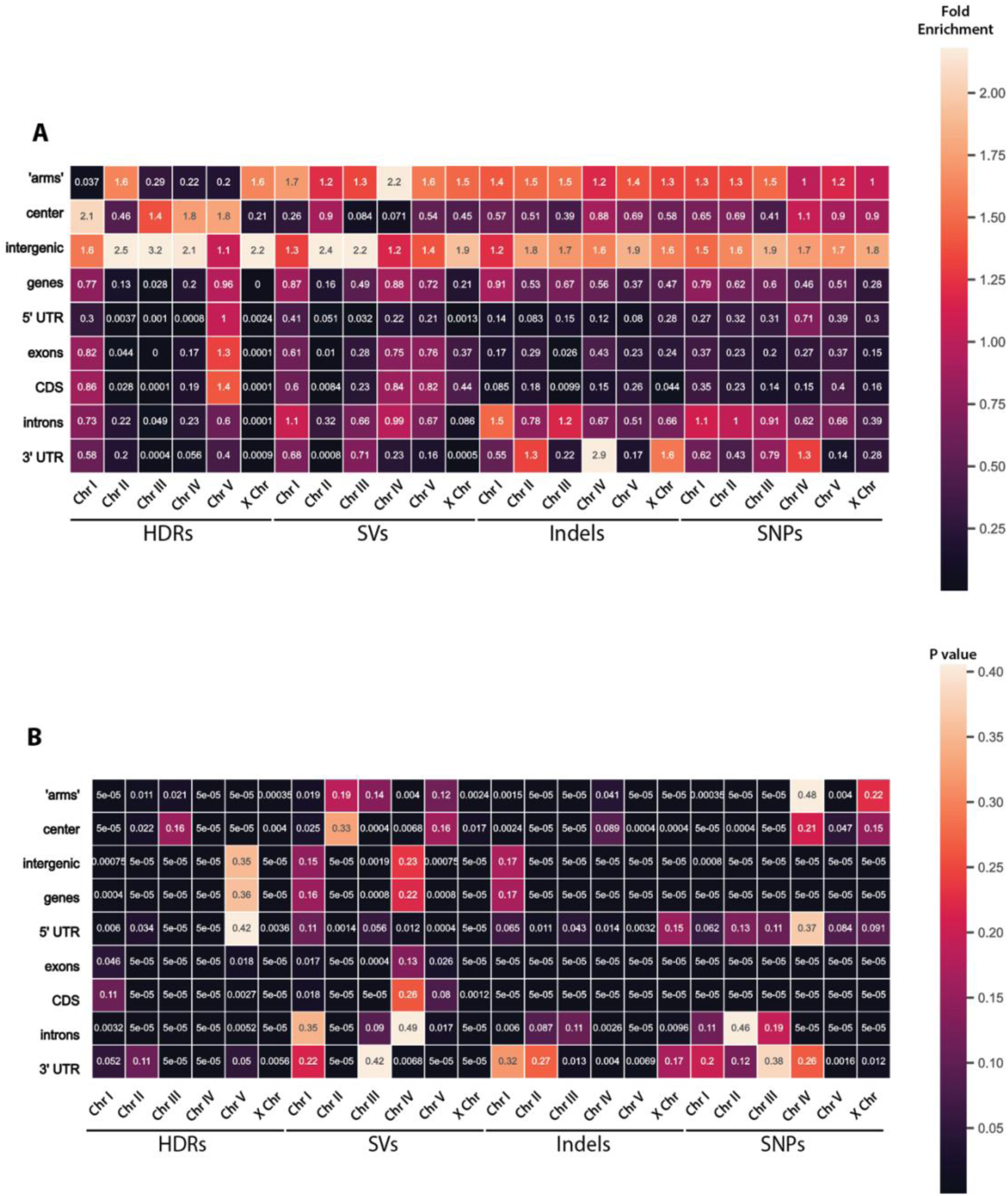
GAT interval-association test results analyzing the overlap of DLW N2 Bristol SNPs, indels, and SVs with remapped VC2010 Bristol genome annotations. A) Heatmap showing the fold enrichment of each variant type within gene annotations for each chromosome. B) Heatmap of p-values associated with corresponding fold enrichments shown in panel A calculated by the hypergeometric test.

**Supplemental Figure S5.**
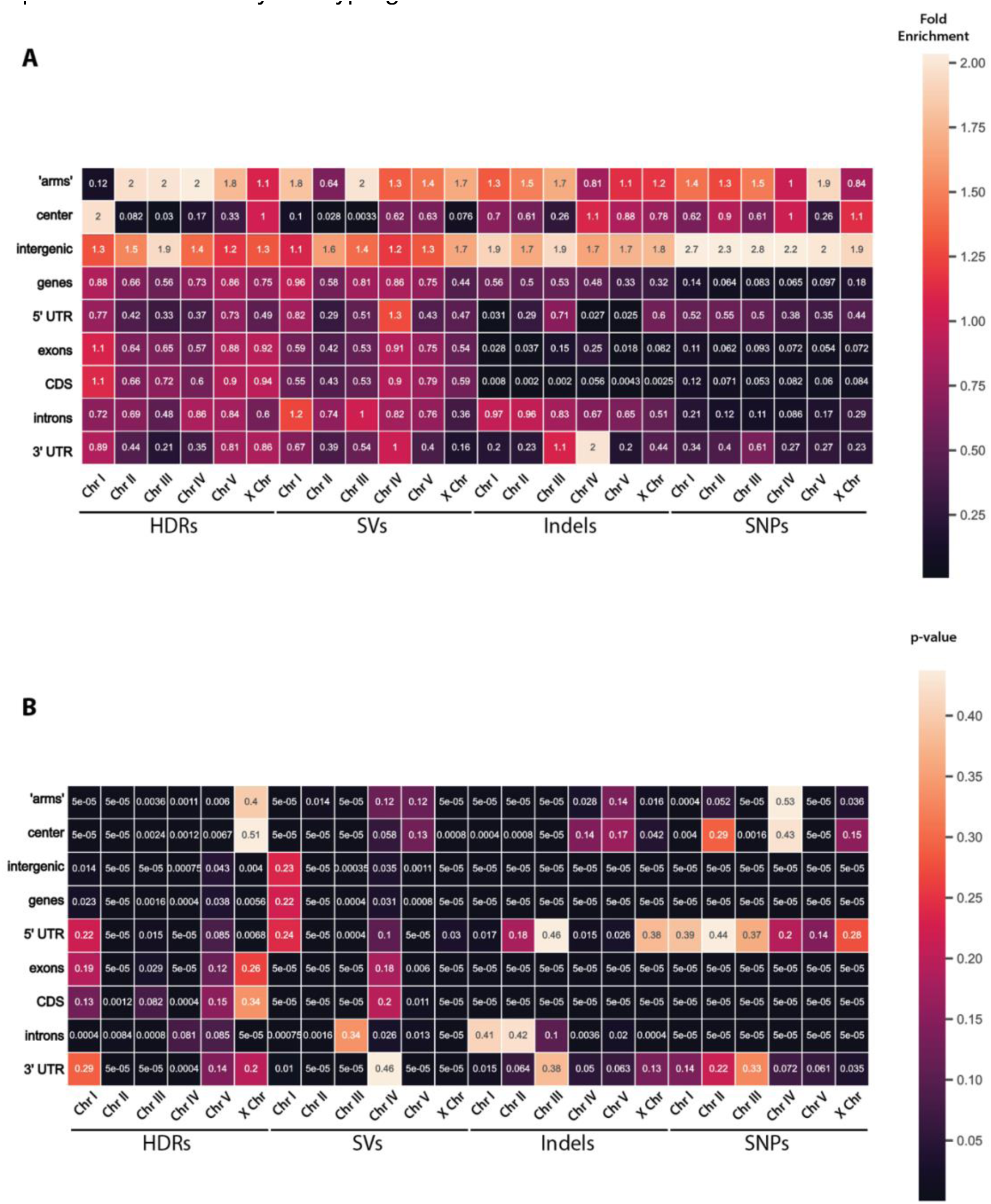
GAT interval-association test results analyzing the overlap of DLW CB4856 Hawaiian SNPs, indels, and SVs with remapped Kim CB4856 Hawaiian genome annotations. A) Heatmap showing the fold enrichment of each variant type within gene annotations for each chromosome. B) Heatmap of p-values associated with corresponding fold enrichments shown in panel A calculated by the hypergeometric test.

**Supplemental Table 1.** Transposable Elements identified in DLW N2 Bristol genome (this study) vs DLW CB4856 Hawaiian genome (this study)

|  | **DLW N2 Bristol** | **DLW CB4856 Hawaiian** |
| --- | --- | --- |
| **Class I Transposable Elements (Retrotransposons)** | 710 (2,688,730 bp) | 776 (2,522,357 bp) |
| **Gypsy** | 557 (2,195,895 bp) | 592 (2,031,038 bp) |
| **Copia** | 134 (472,195 bp) | 161 (465,785 bp) |
| **SINE** | 9 (2,146 bp) | 9 (1,945 bp) |
| **ERV** | 7 (8,280 bp) | 6 (7,569 bp) |
| **LINE** | 3 (10,214 bp) | 8 (16,038 bp) |
| **Class I intrachromosomal transpositions\*** | 0 |  |
| **Class I interchromosomal transpositions\*** | 0 |  |
| **Class II Transposable Elements (DNA transposons)** | 17,682 (12,055,357 bp) | 17,310 (11,606,010 bp) |
| **Tc1/Mariner** | 1870 (1,298,386 bp) | 1,550 (1,131,443 bp) |
| **hAT** | 3,999 (3,988,461 bp) | 3,818 (3,725,667 bp) |
| **CMC** | 1,679 (3,138,647 bp) | 2,011 (3,260,455 bp) |
| **Zator** | 9,159 (3,009,341 bp) | 8,980 (2,907,391 bp) |
| **Novosib** | 46 (12,060 bp) | 28 (12,088 bp) |
| **Helitron** | 39 (368,980 bp) | 43 (329,238 bp) |
| **Sola** | 821 (226,645 bp) | 699 (196,797 bp) |
| **MITE** | 69 (12,837 bp) | 181 (42,931 bp) |
| **Class II intrachromosomal transpositions\*** | 38 |  |
| **Class II interchromosomal transpositions\*** | 9 |  |

\* All TE sequences with predicted transpositions are relative to the DLW N2 Bristol genome

## Notes

https://www.ncbi.nlm.nih.gov/bioproject/907379

### Competing Interest Statement

The authors have declared no competing interest.

### Summary of Updates

New title, abstract, and discussion points to emphasize the novel contributions of the work to our understanding the molecular underpinnings of phenotypic variations and genetic drift of long-term cultivation of laboratory strains

